# Structural Basis for Catalysis and Substrate Specificity of a LarA Racemase with a Broad Substrate Spectrum

**DOI:** 10.1101/2024.11.28.625916

**Authors:** Santhosh Gatreddi, Julian Urdiain-Arraiza, Benoit Desguin, Robert P. Hausinger, Jian Hu

## Abstract

The LarA family consists of diverse racemases/epimerases that interconvert the diastereomers of a variety of α-hydroxyacids by using a nickel-pincer nucleotide (NPN) cofactor. The hidden redox reaction catalyzed by the NPN cofactor makes LarA enzymes attractive engineering targets for applications. However, how a LarA enzyme binds its natural substrate and recognizes different α-hydroxyacids has not been elucidated. Here, we report three high-resolution structures of the enzyme-substrate complexes of a broad-spectrum LarA enzyme from *Isosphaera pallida* (LarA*_Ip_*). The substrate binding mode reveals an optimal orientation and distance between the hydride donor and acceptor, strongly supporting the proposed proton-coupled hydride transfer mechanism. The experimentally solved structures, together with the structural models of other LarA enzymes, allow us to identify the residues/structural elements critically involved in the interactions with different α-hydroxyacid substrates. Collectively, this work provides a critical structural basis for catalysis and substrate recognition of the diverse enzymes in the LarA family, thus building a foundation for enzyme engineering.

## Introduction

The founding member of the LarA family, LarA from *Lactiplantibacillus* (formerly *Lactobacillus*) *plantarum* (LarA*_Lp_*) that interconverts D-/L-lactate,^1, 2^ was established as a nickel-dependent enzyme in 2014^3^ and as a nickel-pincer nucleotide (NPN) cofactor-dependent enzyme in 2015.^4^ The NPN cofactor is synthesized by the sequential reactions catalyzed by three Lar enzymes,^5^ including LarB (a nicotinic acid adenine dinucleotide carboxylase/hydrolase),^5-7^ LarE (an ATP-dependent sulfur transferase),^8-11^ and LarC (a CTP-dependent cyclometallase).^12, 13^ In LarA*_Lp_*, but not all family members, the NPN cofactor is covalently tethered to a universally conserved lysine residue via a thioamide bond.^4^

Accumulating evidence has supported a proton-coupled hydride transfer (PCHT) mechanism for the LarA-catalyzed racemization reaction.^14-17^ According to this hidden redox reaction mechanism, the hydrogen atom on Cα of lactate is transferred as a hydride to the NPN cofactor and then, after a rotation of the acetyl group of the pyruvate intermediate, the hydride returns to Cα to complete racemization. In our previous studies, a strong hydrogen/deuterium primary kinetic isotope effect,^18^ the detection of pyruvate in the reaction mixture,^18^ the reactivity of the NPN cofactor with electron donors, including sulfite and hydride,^18, 19^ and mutagenesis studies^4^ are all consistent with the proposed catalytic mechanism. To elucidate a detailed catalytic mechanism, computational studies have been performed, but only on the putative enzyme-substrate complex models, which are likely to account for different results, including different calculated free energies and even reaction mechanisms.^18, 20-23^ Thus, the lack of an experimentally resolved enzyme-substrate complex structure is a major obstacle to the mechanistic study of LarA enzymes. In our early efforts to resolve the structure of the LarA*_Lp_*-lactate complex, the low substrate affinity, as indicated by the large *K*_M_ values (10-50 mM), and the occupation of the active site by sulfite, a reversible competitive inhibitor and a reductant required to protect the NPN cofactor from rapid oxidative degradation,^4, 19^ prevented us from solving the structure of LarA*_Lp_* in the lactate-bound state.

Our recent bioinformatics study showed that the LarA family consists of distinct subgroups with different substrate specificities.^24^ Sequence analysis of 354 LarA homologs (LarAHs) led to the identification of 25 subgroups, and functional characterization of selected LarAHs revealed that a variety of α-hydroxyacids can be processed by LarAHs, establishing LarA as a racemase/epimerase family. Exchanging the residues in the active sites between LarA*_Lp_* and a LarAH from *Thermoanaerobacterium thermosaccharolyticum*, which processes malate but not lactate, led to swapped substrate preference,^25^ but the structural basis of the substrate specificity has not been established.

In this work, we report the high-resolution crystal structures of LarA from *Isosphaera pallida* (LarA*_Ip_*) in complex with three short-chain aliphatic D-α-hydroxyacids. The structures not only lead to an updated PCHT mechanism but also provide a structural basis of the substrate specificity of α-hydroxyacid racemases/epimerases, paving the way for the identification of the LarAHs catalyzing novel reactions and the rational engineering of LarAHs for potential applications.

## Results

### LarA_Ip_ is an NPN cofactor-dependent enzyme with a broad substrate spectrum

In our previous screen of LarAHs, LarA*_Ip_* (the former LarAH2 or SAR, an abbreviation for short-chain aliphatic α-hydroxyacid racemase) stood out as a promising target for structural studies because of its high affinity for substrates (the *K*_M_ values are 0.15 mM and 0.56 mM for L-lactate and D-lactate, respectively) and its broader substrate spectrum than LarA*_Lp_*.^24^ To prepare the samples in the NPN-bound state for structural studies, the gene encoding LarA*_Ip_* was inserted into an expression vector to allow co-expression in *Lactococcus lactis* with other Lar proteins (LarB, LarC, and LarE) required for the biosynthesis of the NPN cofactor.

A mass spectrometry experiment showed that the purified LarA*_Ip_* has a molecular weight of 47853.5 Da, which corresponds to the protein (47403 Da, after loss of the first methionine) plus 450.5 Da, indicating a covalently tethered NPN cofactor (**Figure 1**). The UV-visible spectrum of the yellow-colored protein showed a broad absorption at 440 nm, which is also present in LarA*_Lp_*. However, the spectrum of LarA*_Ip_* lacks the absorptions at 380 nm and 550 nm seen in LarA*_Lp_*,^18, 19^ suggesting that the NPN cofactor in LarA*_Ip_* is in a different functional state than in LarA*_Lp_*. Compared to LarA*_Lp_*, LarA*_Ip_* is more stable to oxygen in that sulfite is not required to retain functional activity after purification and the active enzyme can be maintained for days.

**Figure 1.**
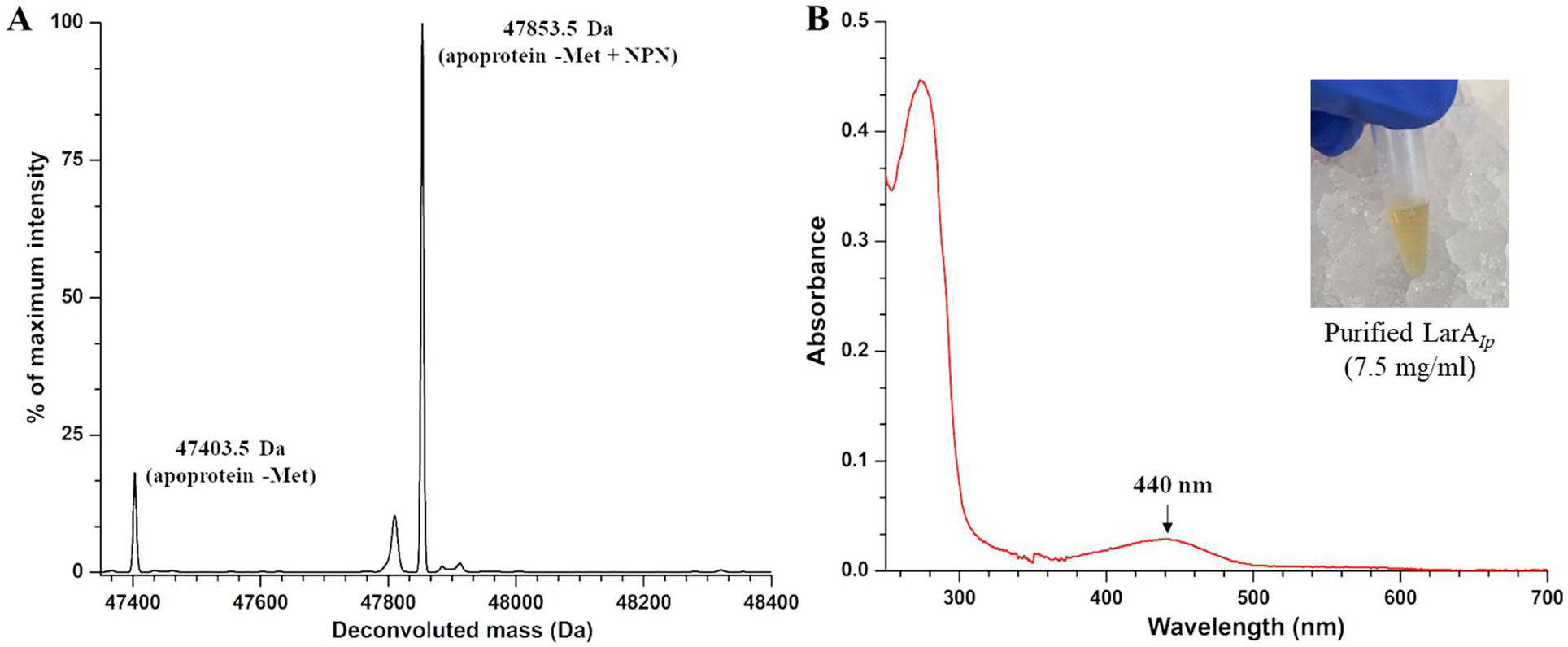
Characterization of the purified LarA*_Ip_*. (**A**) ESI-MS of LarA*_Ip_*. (**B**) UV-visible spectrum of LarA*_Ip_*. The inset shows the appearance of the purified protein.

Extending the early study of this enzyme,^24^ we thoroughly screened an array of α-hydroxyacids and found that LarA*_Ip_* exhibits a broader substrate spectrum than initially reported. As shown in **Figure 2** and **Table S1**, LarA*_Ip_* can process 14 tested compounds with the highest *k*_cat_/*K*_M_ values for L-lactate and 2-hydroxybutyrate (2HB), followed by L-glycerate, 2,4-dihydroxybutyrate, 2-hydroxyvalerate, and 2-hydroxyisovalerate (2HIV). For the compounds with a bulky substituent on Cα, LarA*_Ip_* can still process them but with a much lower activity. For the Cα substituents with similar size, a polar group reduces the reactivity. The broad substrate spectrum of LarA*_Ip_* indicates a larger active site with a greater plasticity than for LarA*_Lp_*.

**Figure 2.**
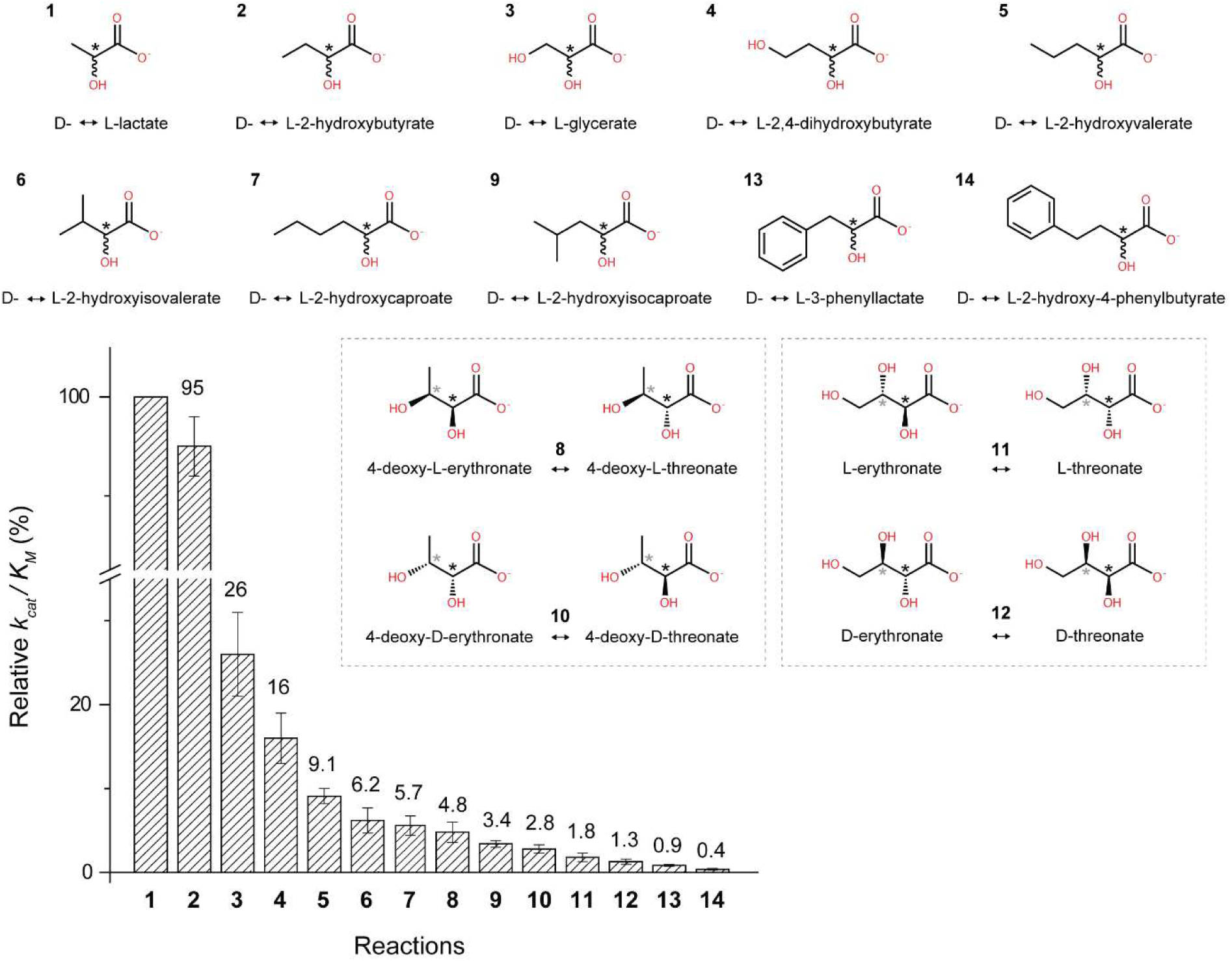
The relative *k*_cat_/*K*_M_ values of LarA*_Ip_* for racemization/epimerization of α-hydroxyacids. The *k*_cat_/*K*_M_ value for lactate was set to 100%. For racemization reactions, the chiral carbons of enantiomers are depicted achiral for simplification. For epimerization reactions, both epimers are depicted, with diastereomers of the same molecule boxed together. Black asterisks indicate stereoinversion sites, grey asterisks indicate other stereocenters in the molecule. The error bars indicate standard deviations (n=3).

### The structure of LarA_Ip_ as purified reveals a naturally bound substrate

We then crystallized LarA*_Ip_* as purified and solved the structure at 1.74 Å (**Table S2**). Two protein molecules are present in one asymmetric unit, but the very small interface (386.2 Å^2^) does not support a physiological dimer. LarA*_Ip_* shares 34.1% identical residues with LarA*_Lp_* (**Figure S1**) and exhibits the same fold (**Figure 3A**). The well-resolved electron density map at the active site clearly indicated a NPN cofactor tethered to Lys183 through a thioamide bond. Ni is coordinated by two sulfur atoms and C4 of pyridinium ring of the NPN cofactor and the fourth ligand is the Nε of His199, completing a nearly ideal square-planar coordination. The NPN cofactor in LarA*_Ip_* occupies the same position and exhibits the identical conformation to that in LarA*_Lp_*, except that the phosphate group of the NPN cofactor adopts a slightly different orientation. Another difference is that the imidazole ring of the side chain of His199 is nearly coplanar with the pyridinium ring of the NPN cofactor in LarA*_Ip_*, whereas in LarA*_Lp_* (PDBs 5HUQ and 6C1W) the two planes are angled by about 50 degrees. The significance of this difference is unclear. The structure of LarA*_Ip_* can be better superimposed with the structure of LarA*_Lp_* in the closed state (PDB 6C1W, chain B, Cα RMSD of 1.10 Å, **Figure 3B**) than in the open state (PDB 5HUQ, chain A, Cα RMSD of 2.0 Å). The N-terminal domains of the two proteins are more similar than the C-terminal domains with the Cα RMSD values being 0.80 Å and 1.41 Å, respectively, consistent with greater sequence identity for the N-terminal NPN cofactor-binding domain (residue 1-269, 36.2%) than that for the C-terminal domain (residue 269-430, 30.3%) (**Figure S1**).

**Figure 3.**
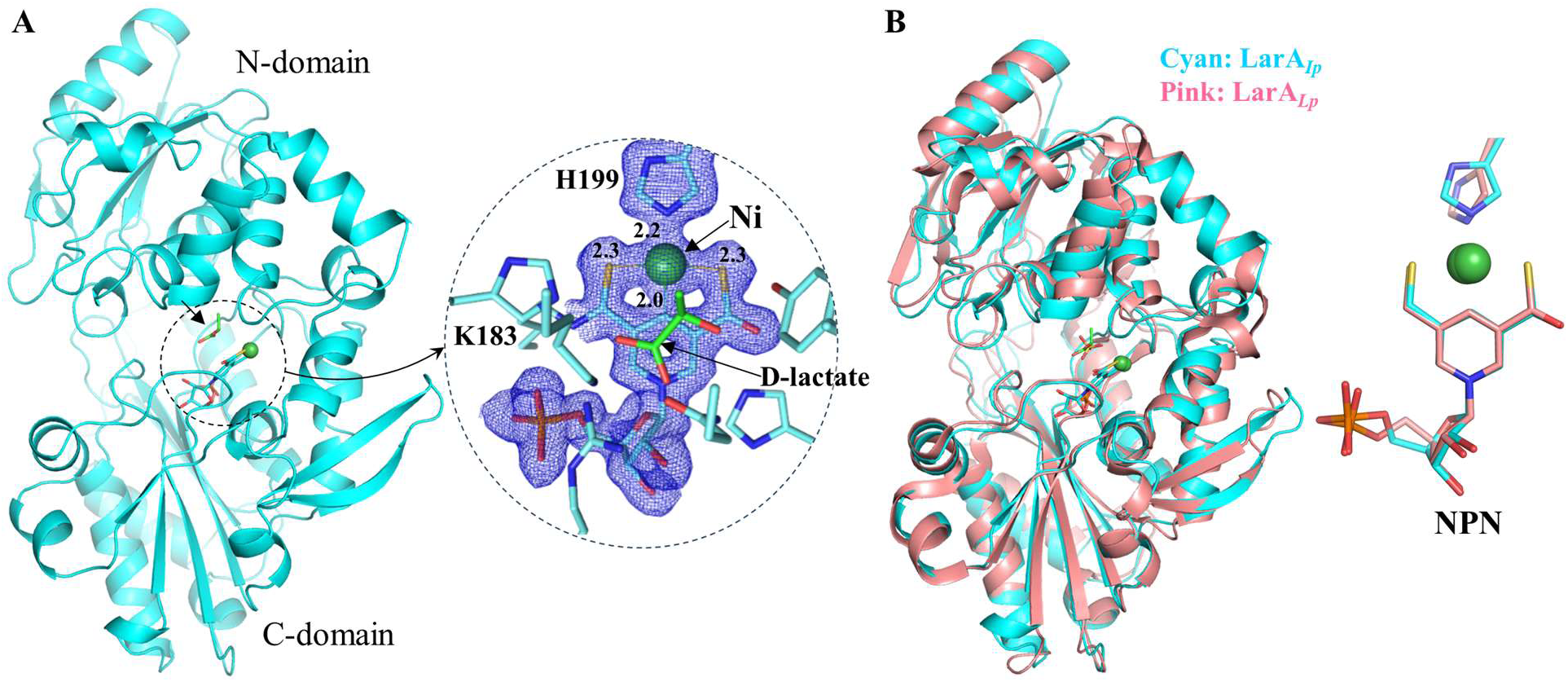
Structure of LarA*_Ip_* as purified. (**A**) Overall structure of LarA*_Ip_* (*left*) and the zoom-in view of the active site (*right*). The blue meshes are the electron densities of the NPN cofactor and His199 (2Fo-Fc, σ=1). (**B**) Structural comparison of LarA*_Ip_* (cyan, chain B) and LarA*_Lp_* (pink, PDB 6C1W, chain B).

On top of the pyridinium ring, a density corresponding to a small molecule was resolved in the structure of LarA*_Ip_* (**Figures 4A** & **S2**). The high quality of the electron density allowed us to assign it as D-lactate (a natural substrate of LarA*_Ip_*) for the best fit of the density. This assignment was later confirmed in a ligand exchange experiment, which is described in a later section. D-lactate is bound at the active site through polar interactions with multiple highly conserved residues (**Figure 4B**, top row). The carboxylic acid group is multi-coordinated with Arg73, Gln295, Lys298, and two catalytic histidine residues His107 and His173, which, together with C2 (or Cα) and the 2-OH group, forms a plane nearly parallel to the pyridinium ring of the NPN cofactor. The rotation of the acetyl group of D-lactate is restricted by the hydrogen bonds between the 2-OH group and residues His107 and Tyr294. Because of these structural constraints, D-lactate is locked in a single conformation with Hα (the hydrogen atom on Cα) pointing toward C4 and Ni of the NPN cofactor with distances of 2.8 Å and 2.9 Å, respectively (**Figure 4B**), allowing a hydride transfer from Cα to the NPN cofactor. Hα is almost identically distant from C4 and Ni, raising a possibility that the hydride can be transferred to Ni rather than C4, as previously proposed for a dual hydride site version of the PCHT mechanism.^18^ These substrate-contacting residues are also conserved in LarA*_Lp_* and involved in the interactions with sulfate at the active site of the enzyme (**Figure 4B**, bottom row).

**Figure 4.**
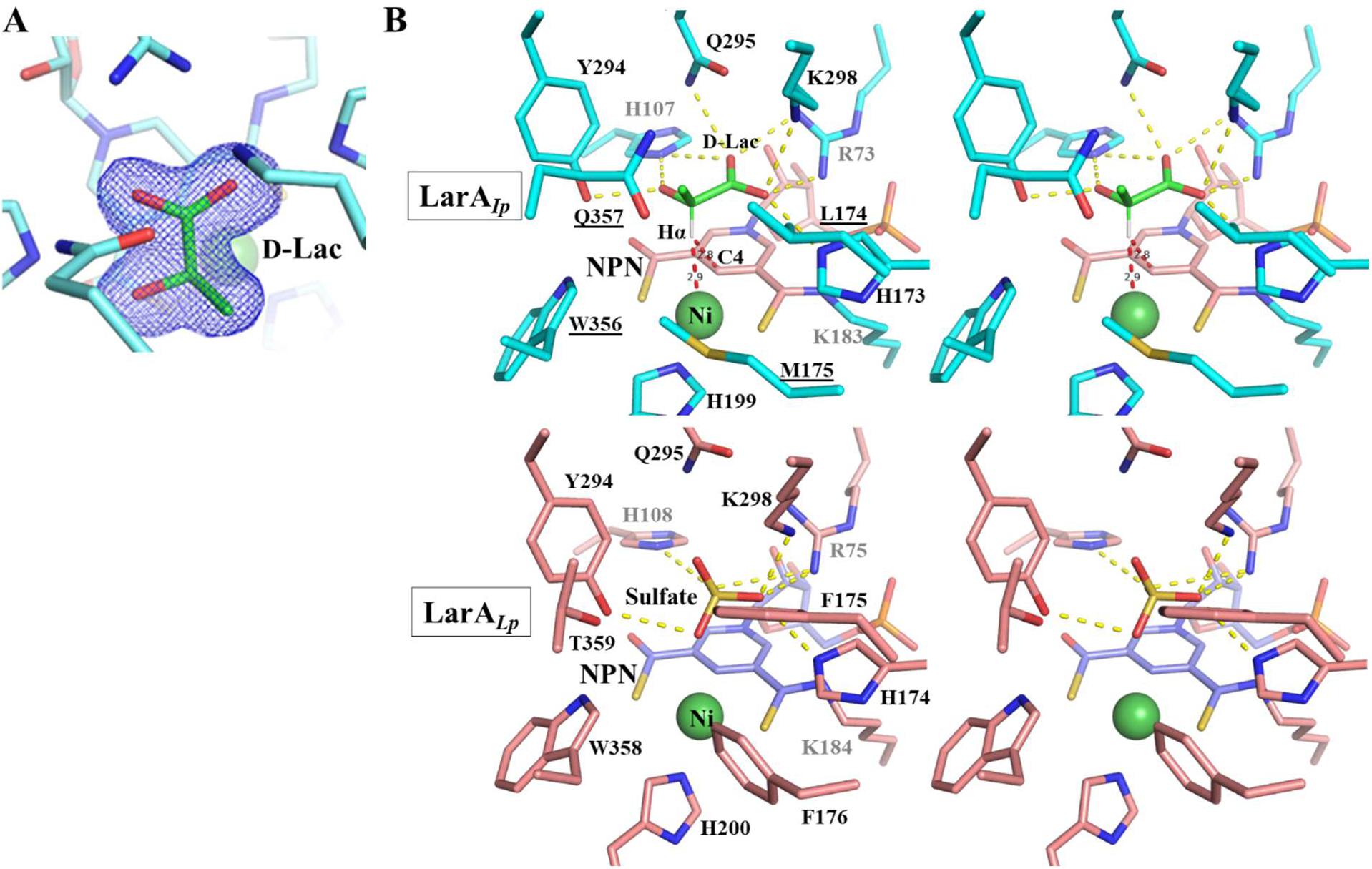
Binding of D-lactate in LarA*_Ip_*. (**A**) 2Fo-Fc map (σ=1) of D-lactate (green) in the active site of LarA*_Ip_* as purified. (**B**) Stereo view of the active sites of LarA*_Ip_* and LarA*_Lp_*. *Top*: LarA*_Ip_* with bound D-lactate. NPN is shown in pink, D-lactate in green, and protein in cyan. The yellow dashed lines indicate polar interactions with D-lactate, and the red dashed lines indicate that the distances between the hydrogen atom on Cα (Hα) and C4 or Ni are 2.8 and 2.9 Å, respectively. The underlined residues are involved in the interactions with the 2-methyl group (Cα substituent). *Bottom*: LarA*_Lp_* with bound sulfate (PDB 6C1W, chain B).

### Co-structures of LarA_Ip_ with additional D-substrates

Given that LarA*_Ip_* has a broader substrate spectrum than LarA*_Lp_* while the active sites of the two enzymes are very similar (**Figure 4B**), we wondered how LarA*_Ip_* recognizes non-lactate substrates. To obtain the enzyme-substrate complexes, we incubated LarA*_Ip_* as purified, which is in the D-lactate bound state, with other known substrates in excess at 4 °C for up to one week before crystallization. However, we only ended up with the same D-lactate bound enzymes in the solved structures, indicating that ligand exchange did not happen. Crystal soaking with ligands did not work either, probably because LarA*_Ip_* was crystallized in the closed conformation where the substrate binding site is not accessible from the water channels in the crystals.

To address this problem, we applied a heating-cooling treatment to promote ligand exchange. *I. pallida* was isolated from a hot spring^26^ and we had previously shown that the enzyme exhibited high activity at 45 °C.^24^ In this work, we further examined the temperature-dependent activity and found that the enzyme exhibited maximal activity at 55-60 °C while barely showing any activity at room temperature (**Figure S3A**), explaining why ligand exchange failed when performed at room temperature. We then examined the heat stability of LarA*_Ip_* by heating the purified protein at 55 °C for 30 min and the profile in size-exclusion chromatography showed that the protein after the heat treatment was monodispersed with a narrow and symmetrical peak (**Figure S3B**), indicating that LarA*_Ip_* is heat stable under these conditions. Based on these results, we applied a heating-cooling treatment to promote ligand exchange for LarA*_Ip_*. In the heating step, the purified protein was incubated with a non-D-lactate substrate at approximately 3 mM at 55 °C for 15 min. In this step, the increased dynamics of the protein at the elevated temperature leads to the opening of the active site to the solvent, allowing the release of the bound D-lactate and the binding of the alternative substrate to the active site. In the cooling step, the temperature was changed rapidly to 4 °C in a thermal cycler, followed by incubation on ice to lock the conformation in the most stable state. Using this protocol, we have successfully solved the crystal structures of LarA*_Ip_* bound with two additional aliphatic α-hydroxyacids, D-2HB and D-2HIV, at a resolution of 1.38 Å and 1.65 Å, respectively (**Figures 5A, B** & **S4**, **Table S2**). It should be noted that D/L-2HB was used in the ligand exchange experiment but only the D-enantiomer was found to be bound in the active site.

**Figure 5.**
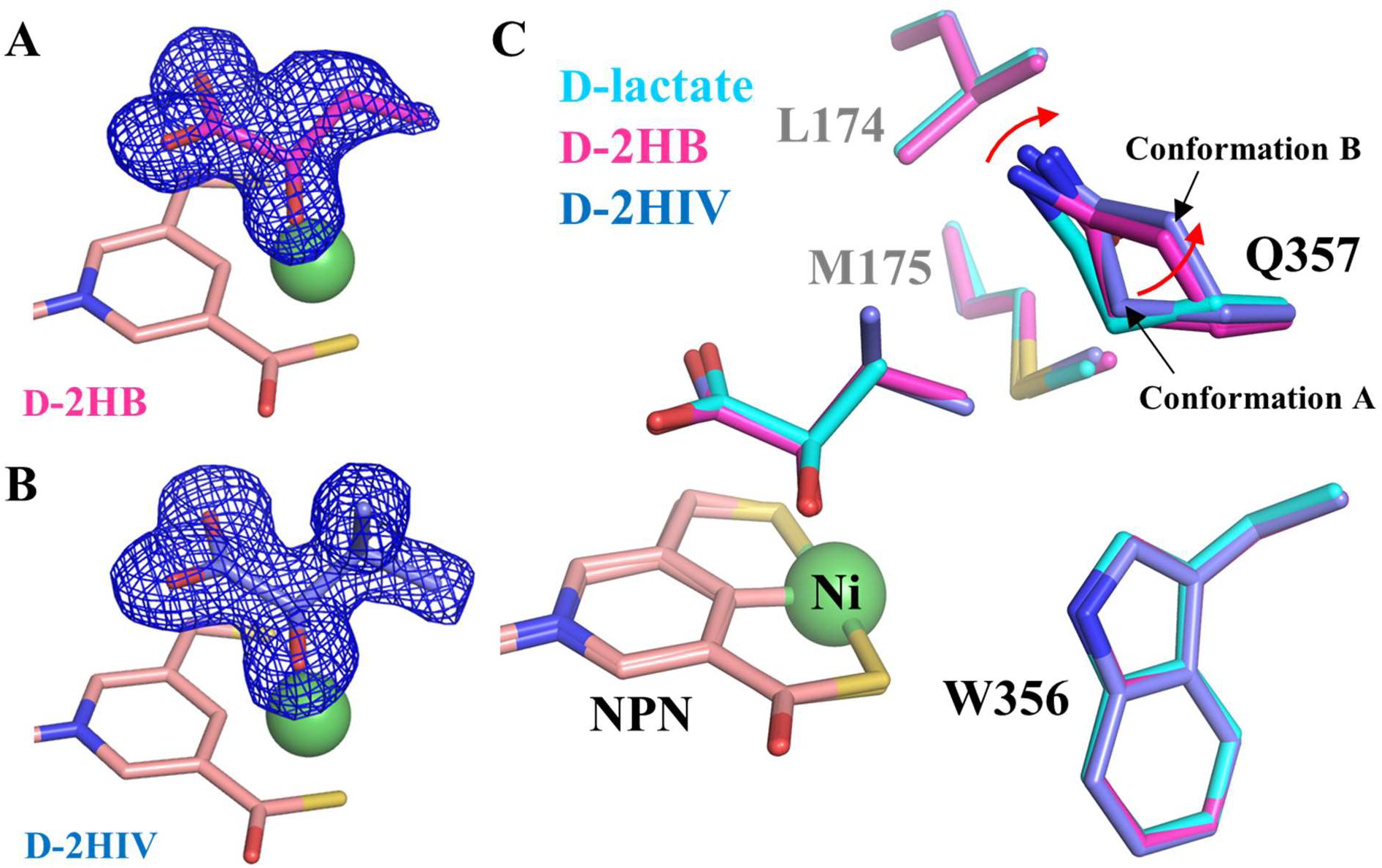
Binding of additional α-hydroxyacids at the active site of LarA*_Ip_*. (**A**) 2Fo-Fc map (σ=1, blue mesh) of D-2HB. The relatively poor density of the terminal methyl group suggests that it is more dynamic than the rest of the ligand. (**B**) 2Fo-Fc map (σ=1) of D-2HIV. (**C**) Structural comparison of the residues contacting the Cα substituents of the different substrates. The red arrow indicates the structural change of Q357 in response to the replacement of D-lactate with 2HB and 2HIV. The NPN cofactor is shown in pink and Ni is shown as a green sphere.

The D-2HB and D-2HIV bound structures can be well superimposed on the D-lactate bound structure with RMSDs of 0.18 Å and 0.13 Å, respectively. The two D-substrates adopt the same orientation as D-lactate and interact with the same set of residues in the active site. Importantly, the aliphatic substituents on Cα of all three substrates point to the same space that is defined by Leu174, Met175, Trp356, and Gln357. Close inspection of these structures revealed that the side chain of Gln357 adopts two conformations (Conformation A and Conformation B) with similar occupancy in the 2HB and 2HIV-bound structures whereas other substrate-contacting residues do not respond to different substrates (**Figure 5C**). In both structures, the amide group in the side chain of Gln357 in Conformation A moves away from the substrate by approximately 0.4 Å, and Conformation B leaves more space to accommodate the substrates with a larger Cα substituent. This result indicates that Gln357 exclusively confers the plasticity of the active site. The geometry and the hydrophobic nature of the substrate binding pocket defined by these residues, as well as the flexibility of Gln357, explained the negative correlation between the size and polarity of the Cα substituents and the reactivity of the substrates (**Figure 2**). In contrast, the bulky and rigid Phe175 and Phe176 in LarA*_Lp_* (**Figure 4B**), which are topologically equivalent to Leu174 and Met175 in LarA*_Ip_*, lead to a smaller and less flexible substrate binding pocket. As a result, LarA*_Lp_* shows a narrower substrate range, whereas LarA*_Ip_* has a broad substrate spectrum with a preference toward those compounds with a small and hydrophobic Cα substituent.

### Heating-cooling treatment with L-substrates

Encouraged by the success of solving the co-structures with D-substrates, we tried to solve the structures of LarA*_Ip_* complexed with L-substrates using the same heating-cooling treatment. Unexpectedly, although the samples treated with L-lactate, L-2HB, and L-2HIV were readily crystallized and the structures were solved at resolutions of 1.44 Å, 1.50 Å, and 1.49 Å (data not shown), respectively, the densities were only consistent with the D-substrates, which must be derived from the added the L-substrates. One possible explanation for observing only the D-enantiomer at the active site, rather than a mixture of enantiomers/anomers as seen in some racemases/epimerases/mutarotases,^27-32^ is that the conformation of LarA*_Ip_* in the L-substrate bound state is different from that in the D-substrate bound state and that the L-substrate bound LarA*_Ip_* was not crystallized in our crystallization trials. Indeed, it is not uncommon that a racemase/epimerase adopts different conformations when bound with different enantiomers.^33^ Alternatively, the D-substrate-enzyme complex has an energy level that is significantly lower than the L-substrate enzyme complex, leading to a preferential binding of the D-enantiomer, as described for alanine racemase.^34^ Although the *K*_M_ value for the L-substrate (0.15 mM) is lower than that for the D-substrate (0.56 mM), *K*_M_ is the sum of *K*_d_ and *k*_cat_/*k*_1_ for an enzyme following the steady-state Michaelis-Menten kinetics; thus, the smaller *k*_cat_ value for the L-substrate than that for the D-substrate, as reported previously,^24^ may be compensated for by the greater *K*_d_ (and thus a lower binding affinity) for the L-enantiomer (**Figure S5**). A similar case was reported in the crystal structure of a variant of a mandelate racemase, where the enantiomer with a smaller *K*_M_ was found to be converted into the enantiomer with a larger *K*_M_.^35^ To reduce the binding of the D-substrates and thus increase the chance of crystallization with the L-substrates, Tyr294, which forms a hydrogen bond with the 2-OH of the D-substrates (**Figures 4B** & **5**), was substituted with alanine. Unfortunately, while the gene encoding the Y294A variant was overexpressed in *L. lactis*, the purified protein did not show any color and the ESI-MS results indicated that the protein did not have a tethered NPN cofactor (**Figure S6**), indicating that Tyr294 plays a role in maintaining the stability of the NPN cofactor. Although we cannot provide structural information of the L-substrate bound enzyme at this point, the ligand exchange experiment crystallographically confirmed the conversion of three L-α-hydroxyacids into the corresponding D-enantiomers, and the resolved L-lactate-derived D-lactate in the active site also unambiguously confirmed the identity of the ligand in LarA*_Ip_* as purified (**Figure 4A**). In addition, the heating-cooling treatment improved the resolution for the D-lactate-bound structure from 1.74 Å to 1.44 Å, likely due to the removal of unstable or poorly folded protein from the sample by the heating treatment.^36^

### Implications of the diverse substrate specificities of LarAHs

The activities and specificities of at least eight subgroups of LarAHs,^24^ including the subgroup containing LarA*_Ip_*, have been experimentally confirmed. Using the binding model for the D-substrates and the AlphaFold predicted structures, we built structural models of enzyme-substrate complexes for the representative members from these subgroups (**Figure 6**). Analysis of these structural models revealed a general model by which LarA enzymes bind different substrates: (1) The N-terminal domain contains three highly conserved residues, including an arginine residue (Arg75 in LarA*_Ip_*) and two catalytic histidine residues (His107 and His173), that bind the carboxylic acid group present in all known LarA enzyme substrates; (2) The two residues immediately following the second catalytic histidine residue (Leu174 and Met175) interact directly with the Cα substituents of the substrates; (3) Additional interactions with substrates come from the residues in the two central α-helices in the C-terminal domain. Some of these residues interact with the carboxylic acid group and 2-OH of the substrates, and the others, together with Leu174 and Met175 (or their equivalents in other LarA enzymes) from the N-terminal domain, define a pocket for recognition of the Cα substituents of the substrates.

**Figure 6.**
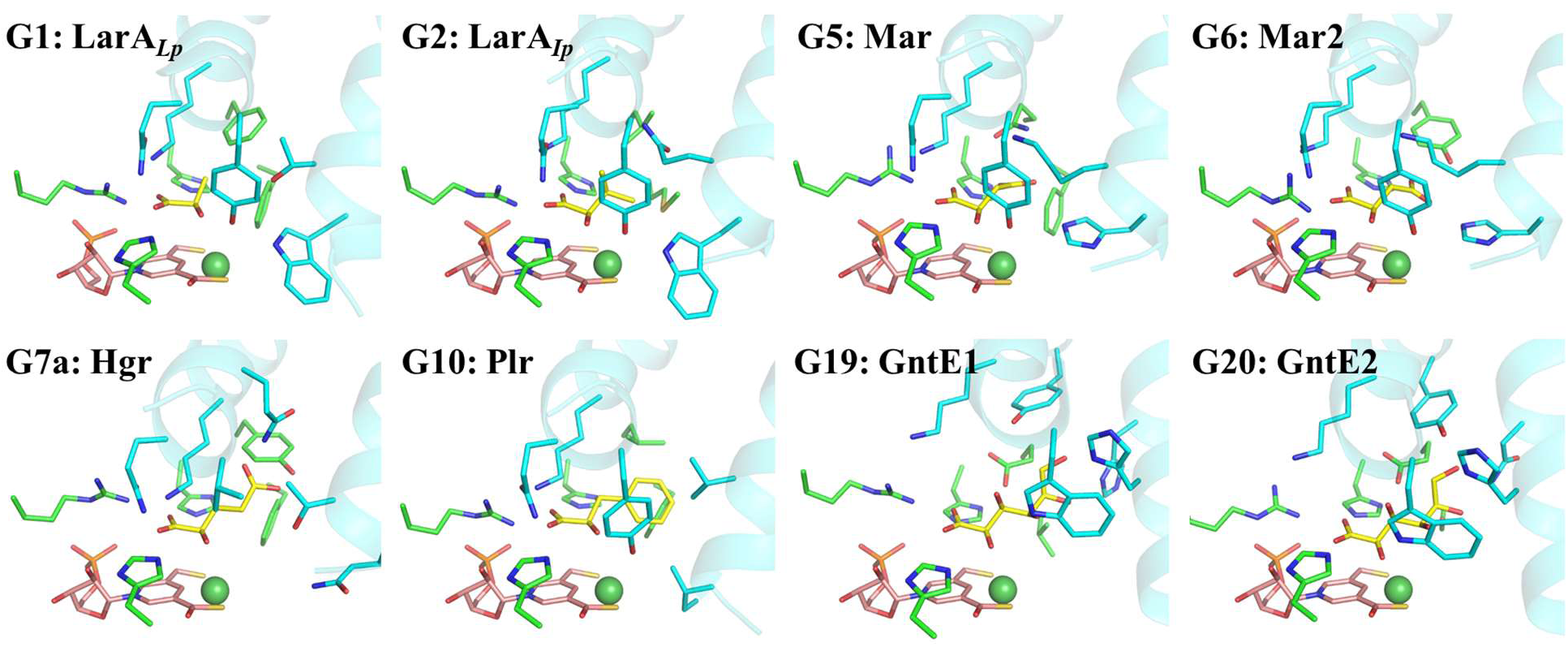
Structural models of the enzyme-substrate complexes for the LarA family members with known substrates. A total of 25 groups of LarA enzymes were identified in a bioinformatics analysis (ref 24), among which eight groups (G1, G2, G5, G6, G7a, G10, G19, and G20) were shown to process at least one α-hydroxyacid. The structures of the enzyme-substrate complexes for representative group members in G5, G6, G7a, G10, G19, and G20 were generated using the AlphaFold predicted models and the D-substrate binding mode revealed in this work. D-lactate was modeled into the structure of LarA*_Lp_* (PDB: 6C1W, chain B) after the removal of the bound sulfate from the active site. G1: lactate racemase; G2: short-chain aliphatic α-hydroxyacid racemase; G5: malate racemase (Mar); G6: malate racemase 2 (Mar2); G7a: hydroxyglutarate racemase (Hgr); G10: phenyllactic acid racemase (Plr); G19: gluconate 2-epimerase 1 (GntE1); G20: gluconate 2-epimerase 2 (GntE2). The substrates are shown in yellow, the NPN cofactors in pink, and Ni in dark green. Only the residues involved in direct interaction with the substrates are shown in stick mode. The residues from the N-terminal domain are in green and those from the C-terminal domain are in cyan. The two central α-helices are shown in cartoon mode.

## Discussion

According to the proposed PCHT mechanism, the NPN cofactor receives a hydride from the substrate and then returns it to the intermediate in one reaction cycle, i.e. a hidden redox mechanism, making LarA enzymes attractive targets for continuous production of desired chemicals as there is no need to supply cofactors during the reactions. An enzyme-substrate complex structure is invaluable in mechanistic studies, including computational studies that rely on a high-quality starting model. This critical missing gap in knowledge is filled by the structures of three D-substrate bound complexes reported in this work. Our data strongly support the PCHT mechanism and also provide key information about the determinants of substrate specificity of LarA enzymes.

The structures reported in this work provide compelling evidence supporting the proposed PCHT mechanism – the 2-OH group of the substrate is H-bonded with the catalytic His107 to allow deprotonation of the former; and Hα faces the NPN cofactor and is within the distance from it as required for a hydride transfer.^37, 38^ Given the nearly identical distance from Hα to C4 or Ni, it is possible that the hydride is transferred to Ni or moves back and forth between C4 and Ni. Since a Ni hydride can be formed in a cationic Ni-phosphine ligand complex with a square pyramidal geometry,^39^ we propose a hypothetical alternative hydride transfer pathway by which the hydride is transferred to Ni with the formed Ni-H bond perpendicular to the distorted planar square (**Figure 7**). Although transient, the formation of such a Ni hydride complex would better allow the rotation of the acetyl group of pyruvate and may also provide an explanation for the selection of Ni for use in the cofactor.

**Figure 7.**
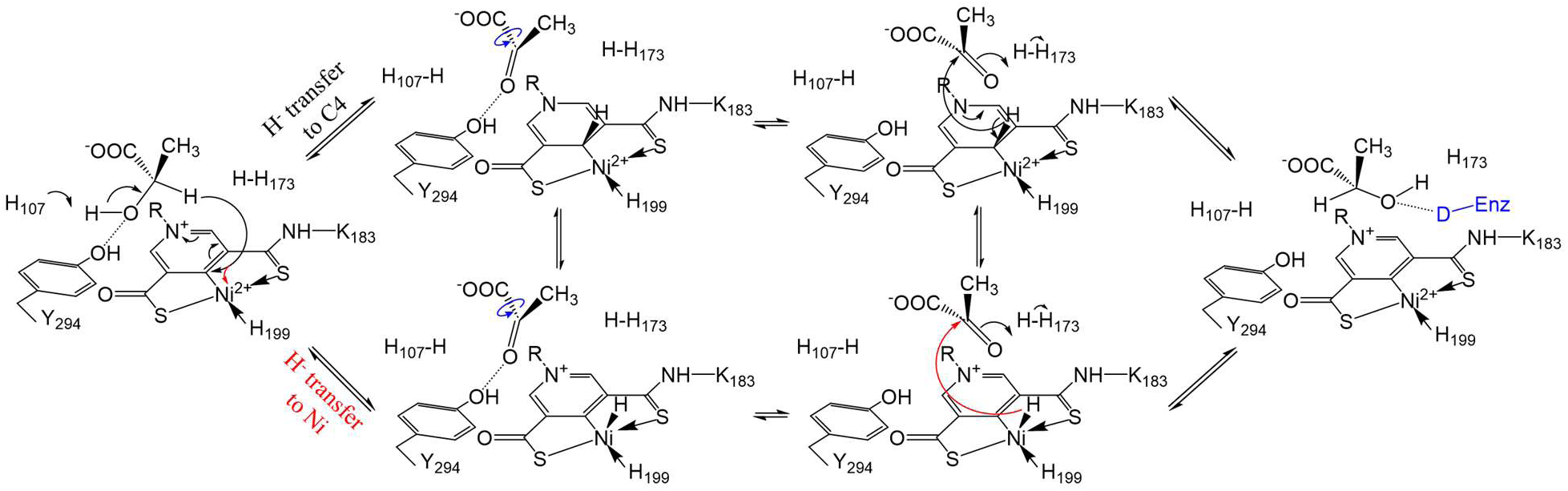
Updated PCHT mechanism of lactate racemization. Two hydride transfer pathways are proposed. The hydride from the D-substrate can be transferred to C4 of the pyridinium ring (black arrow, upper pathway) or to Ni (red arrow, lower pathway). After rotation of the acetyl group of the pyruvate (blue curved arrows) to expose the alternative side of the ketone to the reduced NPN cofactor, the hydride returns to the intermediate to complete racemization. The 2-OH of the product (or L-substrate) is proposed to be stabilized by an unidentified hydrogen donor (capital D in blue), which is equivalent to Y294, in an unresolved conformation of the enzyme.

The solved enzyme-substrate complex structures allowed us to identify multiple residues involved in direct interaction with the substrates. Using the experimentally solved structures and the AlphaFold predicted structures, we have generated additional structural models of the enzyme-substrate complex for the LarA enzymes with confirmed substrates. The compiled structural information leads to a general model for substrate binding to the enzymes. In this model, the N-terminal NPN cofactor-binding domain provides the key contacting residues for the shared carboxylic acid group among the substrates, whereas the C-terminal domain, using the residues on the two central helices, critically contributes to the recognition of different Cα substituents of the substrates (**Figure 6**). In general, the residues involved in binding the carboxylic acid group are more conserved than those interacting with the Cα substituents of the substrates (**Figure S1**). This result is consistent with our previous bioinformatics analysis of 354 LarAHs, showing that the C-terminal domain is more variable than the N-terminal domain.^24^ Given that the LarA family consists of more than 10,000 members according to the InterPro database (LarA_like family, IPR048068), we envision that many additional racemization/epimerization reactions catalyzed by LarA enzymes are certain to be discovered.

In summary, we report the first structures of the enzyme-substrate complex for LarA*_Ip_*, crucially supporting the proposed PCHT mechanism that is likely shared in the entire LarA family and providing the structural basis for substrate specificity. How LarA*_Ip_* or other LarA enzymes bind L-substrates and whether there is a transiently formed high-energy nickel hydride are subjects for future study.

## Materials and Methods

### Cloning, mutagenesis, and protein production/purification

Cloning of the *larAH2* gene encoding LarA*_Ip_* has been described previously.^24^ A vector (pGIR210_LarAH31) for producing LarA*_Ip_* with a C-terminal Strep-II tag in *L. lactis* NZ3900 cells was generated by inserting the *larAH2* gene from the pBADHisA plasmid expressing LarAH2 (processed with restriction enzyme NcoI and NheI) into the plasmid pGIR210 processed with restriction enzymes PciI and NheI.^24^

To generate the Y294A variant, pGIR210 plasmid bearing wild-type LarA*_Ip_* gene was methylated with Dam methylase (New England Biolabs) using S-adenosyl methionine. Methylated template was amplified with the Y294A primers (purchased from Integrated DNA Technologies Inc., **Table S3**) using Phusion^®^ High-Fidelity DNA Polymerase (New England Biolabs). The PCR amplicon was digested with DpnI and desalted for 15 min by drop dialysis using a mixed cellulose ester membrane (0.025 µm size, Merck Millipore Ltd) and then used for transformation into the *L. lactis NZ3900* competent cells by electroporation. Mutation was confirmed by DNA sequencing.

To produce the wild-type and Y294A variant LarA*_Ip_*, the transformed *L. lactis* NZ3900 cells were grown for 14-16 hr at 30 °C in M17 medium supplemented with 0.5% glucose and containing 7.5 μg/mL chloramphenicol. The cultures were then diluted by 100 times with the same medium and incubated at 30 °C with gentle shaking. When the OD_600_ reached 0.3-0.4, 5 μg/L of nisin A and 1mM NiCl_2_ were added and the induction was performed for 3-4 h at 30 °C. The culture was left overnight at 4 °C before the cells were harvested by centrifugation at 4000 x *g* for 15 min. The cells were resuspended in lysis solution containing 100 mM Tris-HCl, pH 7.5, 150 mM NaCl, 2 μg/mL DNaseI, and 10 μg/mL lysozyme, and then stirred at 4 °C for 1 h. After the addition of 1 mM phenylmethylsulfonyl fluoride, cell lysis was performed twice at 16,000 psi using a French press. The supernatant collected after centrifugation at 18,000 x *g* for 70 min was loaded onto StrepTactin XT resin (IBA, Göttingen, Germany), which was pre-equilibrated with 100 mM Tris-HCl, pH 7.5, containing 150 mM NaCl. The resin was extensively washed with the same solution and the Strep-tagged LarA*_Ip_* samples were eluted with 50 mM biotin in the same buffer. The concentrated protein samples were further purified by size-exclusion chromatography on a Superdex 200 increase 10/30 GL column. The fractions corresponding to the protein in the monomeric state were pooled and the concentration was determined by a Nanodrop spectrophotometer using an extinction coefficient of 35,410 M^-1^ cm^-1^.

### UV-Vis spectroscopy

UV-Vis spectrum (250-700 nm) of purified LarA*_Ip_* at a concentration of 7.5 mg/ml in a solution containing 20 mM Tris-HCl, pH 7.4, and 125 mM NaCl was recorded at room temperature on a Shimadzu UV-2600 spectrophotometer (Kyoto, Japan) with 2 nm slit width and 10 mm path length.

### Mass spectrometry

Purified protein (wild-type LarA*_Ip_* or the Y294A variant) at concentrations of 0.4-1 mg/ml in a solution containing 50 mM Tris-HCl, pH 7.4, and 125 mM NaCl was used in the MS experiments. Mass spectra were collected on Xevo G2-XS QTof (Waters) connected to Thermo Hypersil Gold CN guard desalting column (1.0 x 10 mm) after 10 µl protein sample was injected at a flow rate of 0.1 ml/min. The solvents 0.1% formic acid in water (solvent A) and acetonitrile (solvent B) were mixed in a 98%:2% ratio and the percentage of solvent B was gradually increased up to 75%. All data were collected in positive ion mode. Molecular mass spectra were produced with MaxEnt1 algorithm, and the data were plotted using OriginPro 8.

### Enzymatic assays

The general protocol for assaying LarA*_Ip_* purified from *E. coli* in the apoprotein state was as follows. 1 μM of *in vitro* synthesized NPN, 30 mM substrate, and 0.2 μM of purified LarA*_Ip_*, were mixed in 100 mM Tris-HCl, pH 8.0, in 50 μl final volume for 20 min at 30 °C. For *K*_M_ measurements, reactions were performed with variable concentrations of substrate, 1 μM of *in vitro* synthesized NPN, 0.1 μM of purified LarAH, and 100 mM Tris-HCl, pH 8.0, in 50 μl final volume for 20 min at 30 °C.

All reactions were then stopped by incubation at 90 °C for 10 min and the products were assayed spectrophotometrically or by capillary electrophoresis.^40^ D-lactate and L-lactate were assayed spectrophotometrically at 340 nm with an Infinite 200 PRO plate reader (Tecan) using D-/L-lactic Acid Assay Kit from Megazyme. All other racemization and C2-epimerization reactions shown in Figure 2 were assayed by capillary electrophoresis.^40^ In brief, reaction mixtures were loaded onto a polyacrylamide-coated capillary of 54/46 cm total/effective length with an internal diameter of 50 μm from Agilent and run on a Capel 105 M from Lumex Instrument at 20 °C using -25 kV. Using a modified partial filling-counter current method with indirect UV detection, high resolution was achieved with vancomycin as a chiral selector added to the background electrolyte composed of 10 mM of benzoic acid/L-histidine at pH 5. Products were detected at 230 nm.

The relative *k*_cat_/*K*_M_ values were determined by the ratios of the reaction rates of two substrates that were mixed at the same initial concentration in the reaction solution and processed by the enzyme simultaneously.^41^ One substrate was used as the reference and the *k*_cat_/*K*_M_ value of another substrate was reported as the percentage of that the reference substrate. The reaction rates were determined either spectrophotometrically or by capillary electrophoresis,^40^ as described above.

The activity of LarA*_Ip_* produced in *L. lactis* was measured in a solution containing 60 mM MOPS, pH 7.4, and 3 mM sodium L-lactate at the indicated temperatures. The reactions were initiated by adding enzyme (0.5 µM) to pre-equilibrated reaction mix for 10 min and then terminated by heating at 90 °C for 10 min. Precipitated protein was separated by centrifugation, and the quantity of D-lactate formed by racemization activity was measured by using the D-/L-lactate assay kit (Megazyme Inc.), as described previously.^4^

### Ligand exchange by the heating-cooling treatment

Ligand exchange by the heating-cooling treatment was performed in a thermal cycler (T100, Biorad) with a two-step heating-cooling procedure. To study heat stability of LarA*_Ip_*, 4 µM enzyme was mixed with 3 mM sodium L-lactate (Alfa Aesar, stocks prepared in gel filtration buffer containing 20 mM Tris-HCl, pH 7.5, and 125 mM NaCl) in PCR tubes on ice and then transferred immediately to the thermal cycler preheated to 45-65 °C. The samples were heat treated for 30 min and then rapidly cooled to 4 °C. Samples were then centrifuged at 12000 rpm at 4 °C for 10 min to remove the precipitated protein. The supernatants were concentrated and then loaded onto a Superdex 200 Increase 10/300 GL column to check protein oligomeric state as an indicator of protein stability. The gel filtration profiles at 45-65 °C indicated 55 °C as the optimum temperature. To prepare samples for crystallization, ligand exchange (at ∼3 mM) was performed at 55 °C for 15 min followed by rapid cooling to 4 °C as described above, except that different ligands were included during the heating-cooling procedure. The monomeric species collected from gel filtration were concentrated and applied to crystallization.

### Crystallization, data collection, processing, and structure determination

The purified monomeric Lar*_Ip_* was concentrated to 19 mg/ml in a buffer containing 50 mM Tris-HCl, pH 7.4, and 125-300 mM NaCl using a 30 kDa cutoff Amicon ultra centrifugal filter. Crystallization screening was performed using commercial kits by hanging vapor diffusion at 21 °C. Crystals of Lar*_Ip_* grew with a reservoir solution containing 0.1 M imidazole/MES monohydrate, pH 6.5, 20 % Poly(ethylene glycol) methyl ether 500 (PEG 500 MME), 20% PEG 20,000, and 120 mM each of ethylene glycols (diethylene glycol, triethylene glycol, tetraethylene glycol and pentaethylene glycol). The D/L-2-HB crystals grew with 0.1 M Bis-Tris, pH 6.0, and 25% PEG 5000, and D-2-HIV crystals grew with a reservoir solution containing 0.1 M imidazole/MES monohydrate, pH 6.5, 20 % PEG 500 MME, 20% PEG 20,000, and 120 mM each of monosaccharides (D-glucose, D-mannose, D-galactose, L-fucose, D-xylose and N-acetyl-D-glucosamine). Crystals typically appeared within 1-2 days and reached maximum size within 3-4 weeks. All crystals were directly flash frozen in liquid nitrogen.

Diffraction data were collected on 21-ID-D beamline at Life Sciences Collaborative Access Team (LS-CAT) of the Advanced Photon Source in the Argonne National Laboratory and on 17-ID-2 (FMX) beamlines at the National Synchrotron Light Source II at Brookhaven National Laboratory. The dataset for the crystals of LarA*_Ip_* as purified was indexed and scaled using HKL2000, whereas all other datasets were indexed, integrated and scaled using XDS via Fast DP. To conduct molecular replacement with Phenix.phaser, the AlphaFold predicted structure of LarA*_Ip_* was divided into the N-terminal domain and the C-terminal domain. The phases were solved initially by performing molecular replacement using each domain separately before they were combined into a single chain for refinement conducted using Phenix.refine. Model building was performed in Coot. Phenix was used to generate the mF_o_–DF_c_ and 2mF_o_–DF_c_ and maps and Pymol was used to generate images. The atomic coordinates were deposited in the PDB. The crystallographic statistics are listed in **Table S2**.

### Generation of the structural models for the enzyme-substrate complexes

To examine the performance of AlphaFold in predicting LarAHs, the predicted structure of LarA*_Ip_* was compared to the experimentally solved structure. Although the RMSD between the full length LarA*_Ip_* and the predicted structures is 1.86 Å, separate alignment of N- and C-terminal domains showed much smaller RMSD values (0.36 Å and 0.38 Å, respectively), indicating that the large RMSD for the full-length protein is due to different orientations between the two domains. After the N- and C-terminal domains were separately aligned with the experimentally solved structure and then fused into a single protein, the resulting model aligned very well with the experimentally solved structure with an RMSD of 0.36 Å. We treated the AlphaFold models of the LarA enzymes in Groups 1/5/6/7a/10^24^ in the same way to generate the models for manual docking of the NPN cofactor and the corresponding substrates. For the LarA enzymes in Groups 19 and 20, as their C-terminal domain cannot be well aligned with that of LarA*_Ip_*, the AlphaFold models were used for ligand modeling without adjustment. To add the NPN cofactor, the N-terminal domains of LarAHs were aligned with the corresponding region of LarA*_Ip_*, and then the NPN cofactor was directly copied to the LarAH structural models. To add the corresponding substrates to the active site, the binding mode of D-lactate in LarA*_Ip_* was used to guide the docking of other σ-hydroxyacids. Minor adjustment of side chain rotamers and conformations of the ligands were conducted to avoid clashes.

## Acknowledgments

We thank the beamline scientists at LS-CAT of APS and FMX of BNL for collecting data and providing assistance with data processing. We thank Dr. Tony Schilmiller in the Mass Spectrometry & Metabolomics Core Facility at MSU for help with the LC-MS experiments. This work is supported by National Institutes of Health GM128959 (to R.P.H. and J.H.), GM128959 (to R.P.H.), GM140931 (to J.H.), and the UCLouvain Fonds Spécial de Recherche (FSR) (to B.D.).

## Author contributions

J. H., R.P.H. and B.D. conceived the project and designed the experiments; S.G., J.U.A. and B.D. conducted the experiments; S.G., J.U.A., B.D., R.P.H. and J.H. analyzed the data and wrote the manuscript.

## Data Availability

The atomic coordinates and structure factors generated in this study have been deposited in the PDB with the accession codes of 9EIA (LarA*_Ip_* with bound D-lactate as purified), 9EID (LarA*_Ip_* with bound D-2HB), and 9EIF (LarA*_Ip_* with bound D-2HIV). The AlphaFold predicted structures were retrieved from the AlphaFold protein structure database (https://alphafold.ebi.ac.uk/).

## References

1. Iino, T.; Uchimura, T.; Komagata, K., The effect of sodium acetate on the growth yield, the production of L- and D-lactic acid, and the activity of some enzymes of the glycolytic pathway of Lactobacillus sakei NRIC 1071T and Lactobacillus plantarum NRIC 1067T. The Journal of General and Applied Microbiology 2002, 48 (2), 91–102.

2. Goffin, P.; Deghorain, M.; Mainardi, J.-L.; Tytgat, I.; Champomier-Vergès, M.-C.; Kleerebezem, M.; Hols, P., Lactate Racemization as a Rescue Pathway for Supplying d-Lactate to the Cell Wall Biosynthesis Machinery in Lactobacillus plantarum. Journal of Bacteriology 2005, 187 (19), 6750–6761.

3. Desguin, B.; Goffin, P.; Viaene, E.; Kleerebezem, M.; Martin-Diaconescu, V.; Maroney, M. J.; Declercq, J. P.; Soumillion, P.; Hols, P., Lactate racemase is a nickel-dependent enzyme activated by a widespread maturation system. Nat Commun 2014, 5, 3615.

4. Desguin, B.; Zhang, T.; Soumillion, P.; Hols, P.; Hu, J.; Hausinger, R. P., A tethered niacin-derived pincer complex with a nickel-carbon bond in lactate racemase. Science 2015, 349 (6243), 66–9.

5. Desguin, B.; Soumillion, P.; Hols, P.; Hausinger, R. P., Nickel-pincer cofactor biosynthesis involves LarB-catalyzed pyridinium carboxylation and LarE-dependent sacrificial sulfur insertion. Proc Natl Acad Sci U S A 2016, 113 (20), 5598–603.

6. Rankin, J. A.; Chatterjee, S.; Tariq, Z.; Lagishetty, S.; Desguin, B.; Hu, J.; Hausinger, R. P., The LarB carboxylase/hydrolase forms a transient cysteinyl-pyridine intermediate during nickel-pincer nucleotide cofactor biosynthesis. Proc Natl Acad Sci U S A 2021, 118 (39).

7. Chatterjee, S.; Nevarez, J. L.; Rankin, J. A.; Hu, J.; Hausinger, R. P., Structure of the LarB– Substrate Complex and Identification of a Reaction Intermediate during Nickel-Pincer Nucleotide Cofactor Biosynthesis. Biochemistry 2023, 62 (21), 3096–3104.

8. Fellner, M.; Desguin, B.; Hausinger, R. P.; Hu, J., Structural insights into the catalytic mechanism of a sacrificial sulfur insertase of the N-type ATP pyrophosphatase family, LarE. Proc Natl Acad Sci U S A 2017, 114 (34), 9074–9079.

9. Chatterjee, S.; Parson, K. F.; Ruotolo, B. T.; McCracken, J.; Hu, J.; Hausinger, R. P., Characterization of a [4Fe-4S]-dependent LarE sulfur insertase that facilitates nickel-pincer nucleotide cofactor biosynthesis in Thermotoga maritima. J Biol Chem 2022, 298 (7), 102131.

10. Fellner, M.; Rankin, J. A.; Desguin, B.; Hu, J.; Hausinger, R. P., Analysis of the Active Site Cysteine Residue of the Sacrificial Sulfur Insertase LarE from Lactobacillus plantarum. Biochemistry 2018, 57 (38), 5513–5523.

11. Zecchin, P.; Pecqueur, L.; Oltmanns, J.; Velours, C.; Schünemann, V.; Fontecave, M.; Golinelli-Pimpaneau, B., Structure-based insights into the mechanism of [4Fe-4S]-dependent sulfur insertase LarE. Protein Science 2024, 33 (2).

12. Turmo, A.; Hu, J.; Hausinger, R. P., Characterization of the nickel-inserting cyclometallase LarC from Moorella thermoacetica and identification of a cytidinylylated reaction intermediate. Metallomics 2022, 14 (3).

13. Desguin, B.; Fellner, M.; Riant, O.; Hu, J.; Hausinger, R. P.; Hols, P.; Soumillion, P., Biosynthesis of the nickel-pincer nucleotide cofactor of lactate racemase requires a CTP-dependent cyclometallase. Journal of Biological Chemistry 2018, 293 (32), 12303–12317.

14. Hausinger, R. P.; Desguin, B.; Fellner, M.; Rankin, J. A.; Hu, J., Nickel–pincer nucleotide cofactor. Current Opinion in Chemical Biology 2018, 47, 18–23.

15. Nevarez, J. L.; Turmo, A.; Hu, J.; Hausinger, R. P., Biological Pincer Complexes. ChemCatChem 2020, 12 (17), 4242–4254.

16. Chatterjee, S.; Gatreddi, S.; Gupta, S.; Nevarez, J. L.; Rankin, J. A.; Turmo, A.; Hu, J.; Hausinger, R. P., Unveiling the mechanisms and biosynthesis of a novel nickel-pincer enzyme. Biochemical Society Transactions 2022, 50 (4), 1187–1196.

17. Hausinger, R. P.; Hu, J.; Desguin, B., Methods Enzymol, 68:341–371.

18. Rankin, J. A.; Mauban, R. C.; Fellner, M.; Desguin, B.; McCracken, J.; Hu, J.; Varganov, S. A.; Hausinger, R. P., Lactate racemase nickel-pincer cofactor operates by a proton-coupled hydride transfer mechanism. Biochemistry 2018, 57 (23), 3244–3251.

19. Gatreddi, S.; Sui, D.; Hausinger, R. P.; Hu, J., Irreversible Inactivation of Lactate Racemase by Sodium Borohydride Reveals Reactivity of the Nickel–Pincer Nucleotide Cofactor. ACS Catal 2023, 13 (2), 1441–1448.

20. Yu, M. J.; Chen, S. L., From NAD(+) to Nickel Pincer Complex: A Significant Cofactor Evolution Presented by Lactate Racemase. Chemistry 2017, 23 (31), 7545–7557.

21. Zhang, X.; Chung, L. W., Alternative Mechanistic Strategy for Enzyme Catalysis in a Ni-Dependent Lactate Racemase (LarA): Intermediate Destabilization by the Cofactor. Chemistry 2017, 23 (15), 3623–3630.

22. Qiu, B.; Yang, X., A bio-inspired design and computational prediction of scorpion-like SCS nickel pincer complexes for lactate racemization. Chem Commun (Camb) 2017, 53 (83), 11410–11413.

23. Wang, B.; Shaik, S., The Nickel-Pincer Complex in Lactate Racemase Is an Electron Relay and Sink that acts through Proton-Coupled Electron Transfer. Angew Chem Int Ed Engl 2017, 56 (34), 10098–10102.

24. Desguin, B.; Urdiain-Arraiza, J.; Da Costa, M.; Fellner, M.; Hu, J.; Hausinger, R. P.; Desmet, T.; Hols, P.; Soumillion, P., Uncovering a superfamily of nickel-dependent hydroxyacid racemases and epimerases. Sci Rep 2020, 10 (1), 18123.

25. Gatreddi, S.; Urdiain-Arraiza, J.; Desguin, B.; Hausinger, R. P.; Hu, J., Structural and mutational characterization of a malate racemase from the LarA superfamily. Biometals 2022.

26. Giovannoni, S. J.; Schabtach, E.; Castenholz, R. W., Isosphaera pallida, gen. and comb. nov., a gliding, budding eubacterium from hot springs. Archives of Microbiology 1987, 147 (3), 276–284.

27. Au, K.; Ren, J.; Walter, T. S.; Harlos, K.; Nettleship, J. E.; Owens, R. J.; Stuart, D. I.; Esnouf, R. M., Structures of an alanine racemase fromBacillus anthracis(BA0252) in the presence and absence of (R)-1-aminoethylphosphonic acid (L-Ala-P). Acta Crystallographica Section F Structural Biology and Crystallization Communications 2008, 64 (5), 327–333.

28. Frese, A.; Sutton, P. W.; Turkenburg, J. P.; Grogan, G., Snapshots of the Catalytic Cycle of the Industrial Enzyme α-Amino-ε-Caprolactam Racemase (ACLR) Observed Using X-ray Crystallography. ACS Catalysis 2017, 7 (2), 1045–1048.

29. Major, L. L.; Wolucka, B. A.; Naismith, J. H., Structure and Function of GDP-Mannose-3‘,5‘-Epimerase: An Enzyme which Performs Three Chemical Reactions at the Same Active Site. Journal of the American Chemical Society 2005, 127 (51), 18309-18320.

30. Thoden, J. B.; Holden, H. M., High Resolution X-ray Structure of Galactose Mutarotase from Lactococcus lactis. Journal of Biological Chemistry 2002, 277 (23), 20854–20861.

31. Lee, K.-H.; Ryu, K.-S.; Kim, M.-S.; Suh, H.-Y.; Ku, B.; Song, Y.-L.; Ko, S.; Lee, W.; Oh, B.-H., Crystal Structures and Enzyme Mechanisms of a Dual Fucose Mutarotase/Ribose Pyranase. Journal of Molecular Biology 2009, 391 (1), 178–191.

32. Bearne, S. L., Through the Looking Glass: Chiral Recognition of Substrates and Products at the Active Sites of Racemases and Epimerases. Chemistry – A European Journal 2020, 26 (46), 10367–10390.

33. Lundqvist, T.; Fisher, S. L.; Kern, G.; Folmer, R. H. A.; Xue, Y.; Newton, D. T.; Keating, T. A.; Alm, R. A.; de Jonge, B. L. M., Exploitation of structural and regulatory diversity in glutamate racemases. Nature 2007, 447 (7146), 817–822.

34. Spies, M. A.; Woodward, J. J.; Watnik, M. R.; Toney, M. D., Alanine Racemase Free Energy Profiles from Global Analyses of Progress Curves. Journal of the American Chemical Society 2004, 126 (24), 7464–7475.

35. Kallarakal, A. T.; Mitra, B.; Kozarich, J. W.; Gerlt, J. A.; Clifton, J. R.; Petsko, G. A.; Kenyon, G. L., Mechanism of the Reaction Catalyzed by Mandelate Racemase: Structure and Mechanistic Properties of the K166R Mutant. Biochemistry 2002, 34 (9), 2788–2797.

36. Karlsson, A.; Sauer-Eriksson, A. E., Heating of proteins as a means of improving crystallization: a successful case study on a highly amyloidogenic triple mutant of human transthyretin. Acta Crystallographica Section F Structural Biology and Crystallization Communications 2007, 63 (8), 695–700.

37. Dzierlenga, M. W.; Antoniou, D.; Schwartz, S. D., Another Look at the Mechanisms of Hydride Transfer Enzymes with Quantum and Classical Transition Path Sampling. The Journal of Physical Chemistry Letters 2015, 6 (7), 1177–1181.

38. Bahnson, B. J.; Colby, T. D.; Chin, J. K.; Goldstein, B. M.; Klinman, J. P., A link between protein structure and enzyme catalyzed hydrogen tunneling. Proceedings of the National Academy of Sciences 1997, 94 (24), 12797–12802.

39. Cypcar, A. D.; Kerr, T. A.; Yang, J. Y., Thermochemical Studies of Nickel Hydride Complexes with Cationic Ligands in Aqueous and Organic Solvents. Organometallics 2022, 41 (18), 2605–2611.

40. Urdiain-Arraiza, J.; Desguin, B., Versatile capillary electrophoresis method for the direct chiral separation of aliphatic and aromatic α-hydroxy acids, β-hydroxy acids and polyhydroxy acids using vancomycin as chiral selector. Journal of Chromatography A 2024, 1715.

41. Eisenthal, R.; Danson, M. J.; Hough, D. W., Catalytic efficiency and kcat/KM: a useful comparator? Trends in Biotechnology 2007, 25 (6), 247–249.

