## Supplementary Information for "Structural Basis for Catalysis and Substrate Specificity of a LarA Racemase with a Broad Substrate Spectrum"

**N-domain**

```
F9USS9    --MV--------AIDLPYDKRTITAQIDDENYAGKLVSQAATYHNKLSEQETVEKSLDNP    50
E8QWZ4    ---M--------RVTLDYGKTGLNVDLPDDRTLPPLTIRPAP--PLDDPEAEVVRCLAEP    47
B8FVM0    --MK--------TIELPYGHGTQACLIPDDVDCVYGRLKSV--EPTAEAGEQISAALQNL    48
D9TSN9    MGYK--------EISLKYGKGAVDVKIDENMCTVL-YPEDL--PGVEDPMAEVSRSLKDP    49
D3PA49    MTLVEICLRLGDKMKISYGKGFIDVNINKDYDLYQLKIDSAPLS----GKEILERLDNEP    56
G0VL91    MALK--------EFTFNIGENTQSVMLPEEHIVGVMEGKHV--PAVD-VKQATIDCMRHP    49
R0HTW6    MS-----------------QKTITLPIPEGLA------ESA--ALLGGPTEVLNDEQARQ    35
R4NY70    MTVY------------LEGDPLTEEKIKEGLS------KLV--EDLGKVKK---------    31

F9USS9    IGSDKLEELARGKHNIVIISSDHTRPVP-SHIITPILLRRLRSV-APDARIRILVATGFH    108
E8QWZ4    IGSPPLLDLARGKRSACILVCDITRPVP-NPVLLRPILRTLHAAGLATQDILILVATGLH    106
B8FVM0    IGNINFDK-LRNAKSVAIAVSDMTRPVP-SRLIVEKLLPWLAEFGIHGDQITVLVGGGLH    106
D9TSN9    IGKAPLSDLVKGKKDVVILASDITRPSP-SHILIPPITDELNRAGISDDSIKIVFGLGYH    108
D3PA49    IYSENLTYFIKHARKILFIVPDITRKSG-LQIFIKDLIEKIET---FKKEFSIIFATGTH    112
G0VL91    IGSAPLQEKVQKGDKVCLVVADVTRTWNHSNQFLIYIIDELNLAGIPDDDICIVFAQGSH    109
R0HTW6    FILDEVSKLDIDGKTVCMPIPDGTRSGP-HGLMIQAAYDAIAD---RAKSITILIALGTH    91
R4NY70    ---------------VLVVHTDYTRVDF-THLVAKNLYRFLLER--GLKEFHTLNASGTH    73

F9USS9    RPSTHEELVNKYGED---IVNNE-EIVM-HVS-TDDSSMVKIGQLPSG-----------    150
E8QWZ4    RPSTPAEKVEMLSEE---IARTY-RVED-HYG-TRLEEHTYLGTTPNG-----------    148
B8FVM0    HPATQEEMNYILGEE---LPKKI-KQVL-PHDADDQDCLTFLGTSPLG-----------    149
D9TSN9    RKHTDDEKKTLVGEE---[illegible]NRI-KC[illegible]---DHDIDD-[illegible]YVGTTKRG-----------    147
D3PA49    RKVTDEEKKWILTEE---[illegible]GKTKNG-----------    156
G0VL91    RAQTPEEDVRVCGEE---VIRRI-KTYQ-H-D-CMDPTLVDCGTTKLG-----------    150
R0HTW6    AAMDEPSIAKLLGVPSGTIEERFPKATVLNHDWHNPEAIVSLGTIEAAEISRLTSGLLQD    151
R4NY70    RTMKIEEFEKKLGISRNE-----RRVFFHNHEFFNPEALAFVGTLPAGFVSEMTEGDLEE    128

F9USS9    --GDCIINKV-AAEADLLISEGFIESHFFAGFSGGRKSVLPGIASYKTIMANHSGEFIN-    206
E8QWZ4    --VPAWIDSR-YVQADLKIATGLIEPHLMAGYSGGRKLICPGIAAFETVKLWHGPRFLE-    204
B8FVM0    --TPVYVNQY-FAQADFKIVTGMVDAHQFMGFTAGVKGAVIGLGGRETITGNHVRLFQP-    205
D9TSN9    --TPVEVFRE-VYNADFIIATGNLELHYKAGYSGGHKALLPGVCSKNTIEKNHALMFSE-    203
D3PA49    --TPILLNKA-YLEHDTIIPIASVSYHYFAGFGGGRKMILPGIAARKSALNNHKLVLD-E    212
G0VL91    --TPLKLNKH-VVDADKVIIVDGITTHLFAGYGGGRKLILPGVSGFETIQRNHCHALADE    207
R0HTW6    RDMDVQINKL-VAEADVNLVVGPIFPHEVVGFSGGNKYFFPGCSVHDVIDISHWVGALI-    209
R4NY70    E-IPIKVNRLLFEDFDAIFFINGTVPHESTGFSGGLKIVIPGIASTEVVDTFHWAAVLM-    186

F9USS9    -----SPKARTGNLMHNSIHKDMVYAA-----RTAKLAFIINVVLDEDKKII------GS    250
E8QWZ4    -----HPLADCGFLEGNPVHEENTRIA-----RMAGCDFIVNVTLDGARRIT------SV    248
B8FVM0    -------GAELGQMEGNPARIDLEDCG-----RIIGVDMIVNVVLNTQKKVV------KA    247
D9TSN9    -------GAMPGKIDGNPMREDIEEGG-----KLARVDFIVNAVLNSHKEIV------KV    245
D3PA49    RNMRKHPLATTGNLKQNPVNDDIVEAVM--IARRGKFFFTINTILNDKGEII------DL    264
G0VL91    FGHGINPKTRALKIDDNPLNDDMVEAC-----SKINPCFLVHSVINGDGEIC------RM    256
R0HTW6    -----TASEIIGTLGITPVRQLINSSSALIPGEKLAVTYV----STTGDDDQPVLH--SV    258
R4NY70    -----GIPKLIGTVDN-PARKIINRASEMIFEKIKARSFTLNMVYEEEEEVIPRALYIDE    240

F9USS9    FAGDMEAAHKVGCDFVKELSSV-PAIDCDIAISTNGGYPLDQNIYQAVKGMTAAEAT--N    307
E8QWZ4    VAGDMEQAFLKGVAFVETVVKAAVPAPVDVVVTSSAGHPLDLTFYQAVKGLTGALPI--V    306
B8FVM0    VAGHPRTAHGVAVEFAKSIFGV-PMSSADIVIASPGGFPKDINAYQAQKALTPALQL--V    304
D9TSN9    VSGDPIKAHREGAKYIDKMYKRVIPEKADIVVASCGGYPKDINLYQAQKGLDNAQYS--V    303
D3PA49    TCGDLFMSHIEATERLKKYTMITANKKYDTIIVSCGGYPKDINMVQAQKSLDRVIPI--A    322
G0VL91    VGGDWYDAWRAGTEAVLDIQRVPMKEKADVVIACAGGYPSDVSLYQGCKCYDPAEMA--V    314
R0HTW6    AVGTTESAWAANANVASATHIKWLDAPYKRIVSKIPEMY--EDLWTGAKGVYKMEPV--C    314
R4NY70    GYEGFLRAYEKACELSSQLHVKYIDRPLRRAVQVIGEEY--DEVWTAGKGSYKLQRPGVM    298

F9USS9    KEGGTIIMVA-GARDGHGGEGFYHNLADVDD-PKEFLDQAINTPRLKTIPDQWTAQIFAR    365
E8QWZ4    KPGGTIVIAA-ALAEGLGSPEFQSLFEEHPT-LEGFMEAILKEE--SFTVDQWQLEELAK    362
B8FVM0    KPGGVIILVA-QCSEGSGEESFAKTMALYDN-PSDLVTSFKEKE--FVIG-PHKAYLWTR    359
D9TSN9    KDGGTIILVA-ECREGLGEKLFSDWMVNSSS--VDEPLKWIKEE--FRLG-AHKAAVICE    357
D3PA49    ANNANIIFFA-ECVDGYGNNYFEEFFDITTS--EEMFKT-LIKD---YQINRQTAYSLKI    375
G0VL91    KDGGVIIAIM-EARDIKEPAIYMDSFKY-DT--MEEMETALRAH---FTIPFFVAFNLFC    368
R0HTW6    TDGGEVIVYAPHITEIS---EMHQGLADIGYHCIEYFTKQWDKF-----KDHPWGEIAHS    360
R4NY70    AKGGQIIIYAPHIKRFHSN[illegible]DKWI[illegible]EIGYHCKDYVK[illegible]YLKKH-----PDFNKNVAAHV    347
```

**C-domain**

```
F9USS9    ILVHH-------------HVIFVSDLVDPDLITNMHMELAKTLDEAMEKAYA-----RE    406
E8QWZ4    VRRKA-------------RVKFVSDGVPAAVLSRCHVEPVATVELAVAQALE-----QY    403
B8FVM0    TFLKA-------------KTILVSDKVSPELAKALMVKVTKSLQEAIDDVIP-----DD    400
D9TSN9    VLKRA--------------DIYLISSFDRSLTEKIFFKYAKTPQDALDEAIK-----K-    396
D3PA49    KTENY---------------NVFLYSNFSESDCKRMGFIKINSIEEINNIINN------    411
G0VL91    LTHKD-------------TVILVTLPRNFDDIRRTGQIPVATVQEAWDLAQQKLKEQGK    413
R0HTW6    THVRGLGSFDPETGEEKLRINVTLASQVSPEVCAAYNIGYADPASFDWDALDT-------    418
R4NY70    INVRGAGTFDPETGKEEFEFDVILATSIPEDECRAVNLGYMDPSKIKKEDFM--------    405

F9USS9    GQAAKVTVIPDGLGVIVK----------    424
E8QWZ4    GPEARVAVIPKGPYVLPVVDPTLGTAG-    430
B8FVM0    TAGLKITVLPNANSVIPILRDESNETET    428
D9TSN9    YHDPKILVLPYANSTLPYVEE-------    417
D3PA49    NNANNIAIVPDAYNVFFNTD--------    431
G0VL91    DKDYTINIMPHATKVMPILQEK------    435
R0HTW6    -TDPDTLVVEHAGEILHRLANQRSV---    442
R4NY70    -DE-DSLWIVPGGKYLYDLKERRG----    427
```

**Figure S1.** Sequence alignment of LarA enzymes with known substrates. The Uniprot IDs and the corresponding LarA enzymes are listed below. F9USS9 – LarA*Lp* (group 1); E8QWZ – LarA*lp* (group 2); B8FVM0 – Mar (group 5); D9TSN9 – Mar2 (group 6); D3PA49 – Hgr (group 7); G0VL91 – Plr (group 10); R0HTW6 – GntE1 (group 19); R4NY70 – GntE2 (group 20). The colored residues are shown for LarA*lp* or predicted (for all others) to directly interact with D-α-hydroxyacid substrates: red – Cα substituent; green – carboxylic acid group; blue – α-hydroxyl group. The highlighted sequences form two helices that extensively interact with substrates in the C-terminal domain. The N- and C-terminal domains are shown in the light blue and brown boxes, respectively.

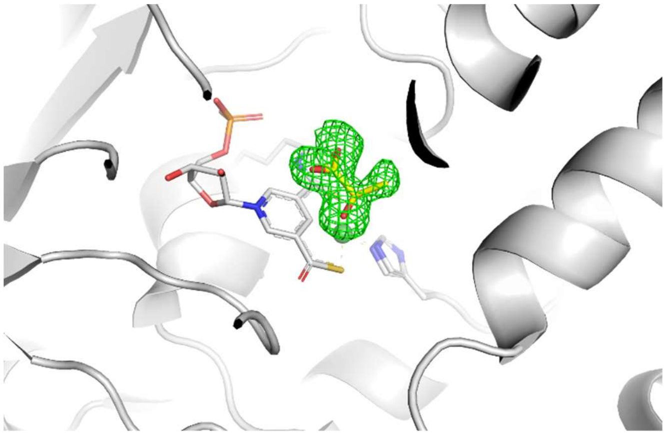

**Figure S2.** Fo-Fc omit map (green meshes, σ=3) of D-lactate (stick mode in yellow) in $LarA_{lp}$ as purified (Chain A). D-lactate, the NPN cofactor, Lys183, and His199 are shown in stick mode.

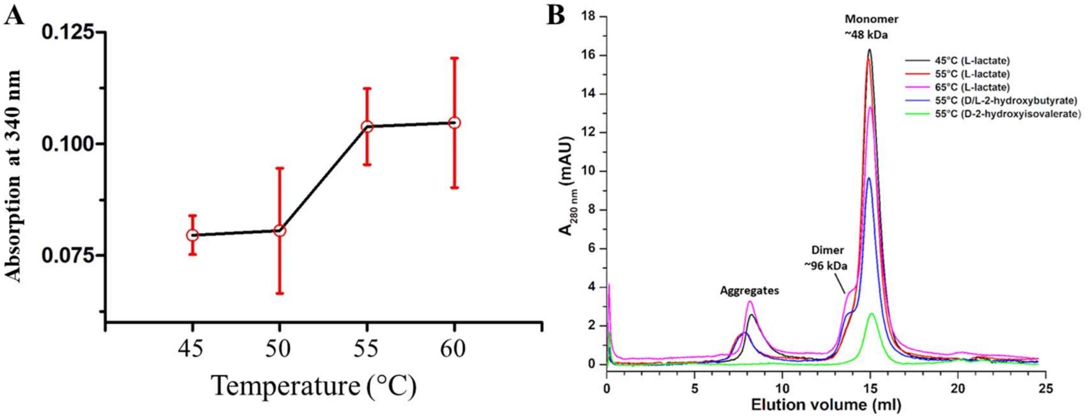


**Figure S3.** Heat stability test of LarA$_{lp}$. (**A**) Temperature dependence of LarA$_{lp}$ activity. L-lactate racemase activity was measured at the indicated temperatures. The error bars indicate standard deviations (n=3). (**B**) Heat stability test of LarA$_{lp}$. Monomeric LarA$_{lp}$ as purified (4 μM) was heated at the indicated temperatures in 20 mM Tris-HCl (pH 7.5) and 125 mM NaCl for 30 min in the presence of 3-5 mM substrates and then cooled rapidly to 4 °C. The stability of the treated samples was evaluated by size-exclusion chromatography.

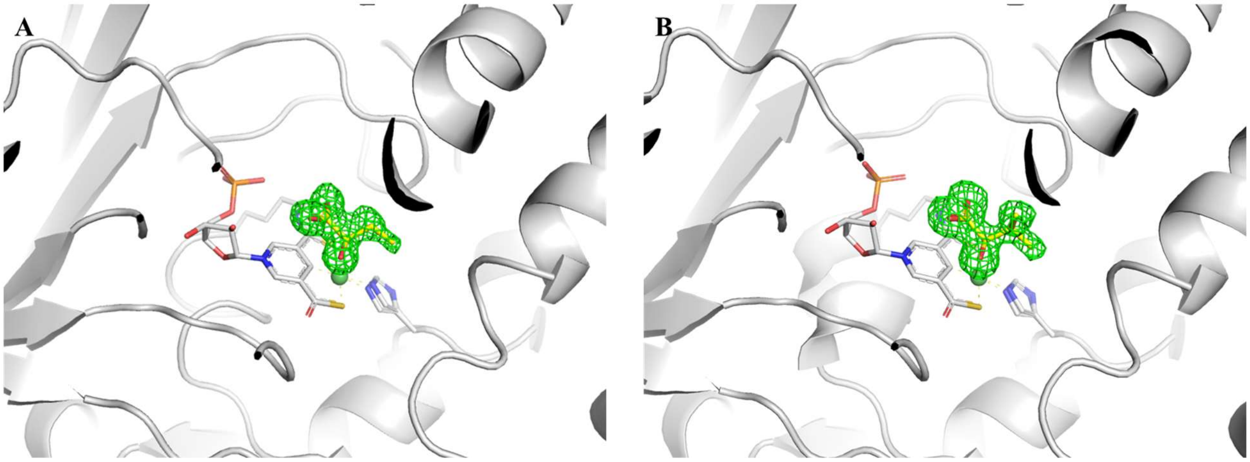


**Figure S4.** Fo-Fc omit maps (green mesh, σ=3) of LarA*lp* after ligand exchange with D-2HB (left) and D-2HIV (right). Substrates (yellow), the NPN cofactor, Lys183, and His199 are shown in stick mode.

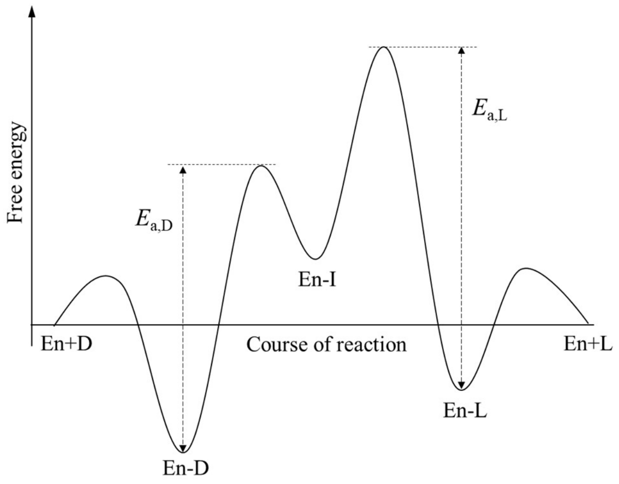


**Figure S5.** Proposed free energy profile of the racemization reaction catalyzed by $LarA_{lp}$. En: enzyme. D: D-enantiomer. L: L-enantiomer. En-D/L: enzyme-substrate complex. En-I: enzyme-intermediate complex. $E_{a,D/L}$: activation energy of the racemization reaction using D- or L-enantiomer as substrate.

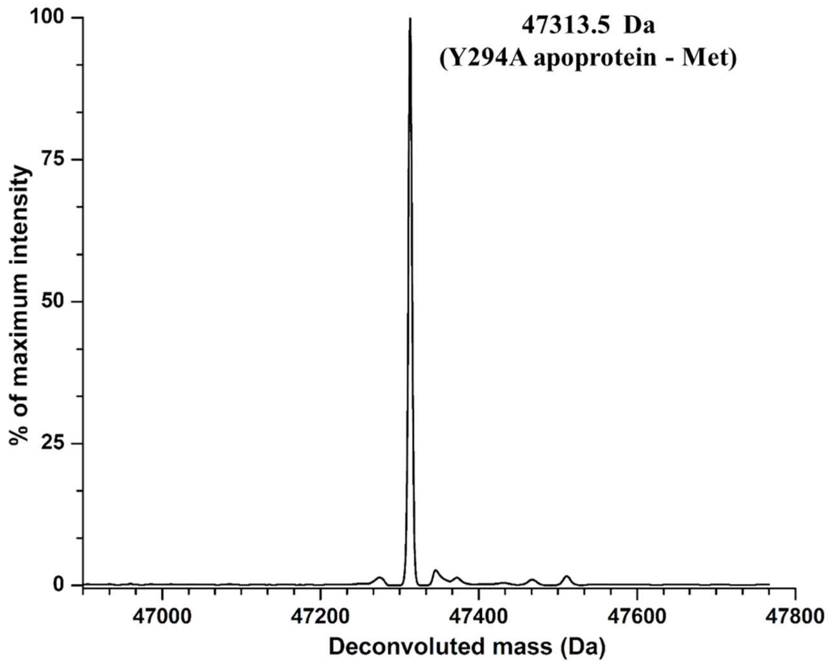


**Figure S6.** ESI-MS of the Y294A variant as purified from *L.lactis*.

**Table S1.** $k_{cat}/K_M$ values of $LarA_{lp}$ for α-hydroxyacids.

| Reactions/Substrates | Relative $k_{cat}/K_M$ (%) | $k_{cat}/K_M$ ($M^{-1}$ $s^{-1}$) |
|---|---|---|
| D- ↔ L-lactate | 100 | (31 ± 9) x $10^3$ |
| D- ↔ L-2-hydroxybutyrate | 95 ± 3 | (30 ± 8) x $10^3$ |
| D- ↔ L-glycerate | 26 ± 5 | (82 ± 27) x $10^2$ |
| D- ↔ L-2,4-dihydroxybutyrate | 16 ± 3 | (51 ± 16) x $10^2$ |
| D- ↔ L-2-hydroxyvalerate | 9.1 ± 0.9 | (28 ± 8) x $10^2$ |
| D- ↔ L-2-hydroxyisovalerate | 6.2 ± 1.5 | (19 ± 7) x $10^2$ |
| D- ↔ L-2-hydroxycaproate | 5.7 ± 1.1 | (18 ± 6) x $10^2$ |
| 4-deoxy-L-threonate ↔ 4-deoxy-L-erythronate | 4.8 ± 1.2 | (15 ± 5) x $10^2$ |
| D- ↔ L-2-hydroxyisocaproate | 3.4 ± 0.4 | (11 ± 3) x $10^2$ |
| 4-deoxy- D-threonate ↔ 4-deoxy-D-erythronate | 2.8 ± 0.5 | (89 ± 28) x $10^1$ |
| D-threonate ↔ L-erythronate | 1.8 ± 0.4 | (56 ± 20) x $10^1$ |
| D-threonate ↔ D-erythronate | 1.3 ± 0.3 | (42 ± 14) x $10^1$ |
| D- ↔ L-3-phenyllactate | 0.85 ± 0.09 | (27 ± 8) x $10^1$ |
| D- ↔ L-2-hydroxy-4-phenylbutyrate | 0.39 ± 0.08 | (13 ± 4) x $10^1$ |

**Table S2.** Crystallographic statistics.

| **Data collection** | **LarA$_{lp}$ as purified (D-lactate)** | **D-2-hydroxybutyrate** | **D-2-hydroxyisovalerate** |
|---|---|---|---|
| Beamline | LS-CAT 21-ID-D | NSLSII 17-ID-1 FMX | NSLSII 17-ID-1 FMX |
| Wavelength (Å) | 1.127231 | 0.97934 | 0.97934 |
| Space group | $P2_1$ | $P2_12_12_1$ | $P2_1$ |
| Unit cell a, b, c (Å); α, β, γ (°) | 79.42, 45.51, 119.14<br>90.00, 91.20, 90.00 | 46.72, 79.61, 104.78<br>90.00, 90.00, 90.00 | 79.02 45.25 118.22<br>90.00 91.07 90.00 |
| [a]Resolution (Å) | 33.07 – 1.74 (1.80-1.74) | 29.11 – 1.38 (1.41-1.38) | 29.71 – 1.65 (1.68-1.65) |
| [a]Redundancy | 2.5 (2.0) | 12.8 (7.8) | 6.9 (6.9) |
| [a]Completeness (%) | 96.2 (94.3) | 99.7 (95.7) | 99.1 (93.4) |
| [a]$I/\sigma I$ | 10.3 (1.52) | 18.6 (2.9) | 10.9 (2.6) |
| [a,b]$R_{merge}$ | 0.112 (0.567) | 0.095 (0.763) | 0.118 (0.716) |
| [a,c]$R_{pim}$ | 0.083 (0.469) | 0.027 (0.288) | 0.048 (0.291) |
| [d]$CC_{1/2}$ of the highest resolution shell | 0.610 | 0.805 | 0.762 |
| **Refinement** | | | |
| Unique reflections | 84,457 | 80,218 | 100,330 |
| Number of atoms | 6965 | 3851 | 7156 |
| Protein atoms | 6339 | 3280 | 6429 |
| $H_2O$ molecules | 519 | 529 | 637 |
| Phosphate | 0 | 2 | 0 |
| EDO | 3 | 0 | 0 |
| PEG | 3 | 0 | 2 |
| PGE | 0 | 0 | 1 |
| Substrate (D-lactate/D-2-hydroxybutyrate/D-2-isovalerate) | 2 | 1 | 2 |
| Ni | 2 | 1 | 2 |
| 4EY | 2 | 1 | 2 |
| [e]$R_{work}/R_{free}$ | 0.174/0.212 | 0.151/0.166 | 0.161/0.188 |
| *B*-factors (Å$^2$) | 18.5 | 13.8 | 17.3 |
| Protein atoms | 17.9 | 12.2 | 16.5 |
| $H_2O$ molecules | 26.5 | 23.5 | 26.1 |
| Phosphate | - | 21.4 | - |
| EDO | 34.0 | - | - |
| PEG | 36.8 | - | 36.1 |
| PGE | - | - | 32.3 |
| Substrate (D-lactate/D-2-hydroxybutyrate/D-2-isovalerate) | 15.3 | 12.5 | 12.7 |
| Ni atoms | 12.3 | 7.4 | 11.7 |
| 4EY: P2TMN | 11.1 | 7.6 | 10.8 |
| R.m.s. deviation in bond lengths (Å) | 0.007 | 0.006 | 0.007 |
| R.m.s. deviation in bond angles (°) | 0.983 | 0.980 | 1.03 |
| Ramachandran plot (%) favored | 98.5 | 98.4 | 98.3 |
| Ramachandran plot (%) allowed | 1.5 | 1.6 | 1.7 |
| Ramachandran plot (%) outliers | 0 | 0 | 0 |
| Rotamer (%) outliers | 0 | 0 | 0 |
| PDB ID | 9EIA | 9EID | 9EIF |

[a]Highest resolution shell is shown in parentheses.
[b]$R_{merge} = \sum_{hkl} \sum_j |I_j(hkl) - \langle I(hkl) \rangle| / \sum_{hkl} \sum_j I_j(hkl)$, where $I$ is the intensity of reflection.
[c]$R_{pim} = \sum_{hkl} [1/(N-1)]^{1/2} \sum_j |I_j(hkl) - \langle I(hkl) \rangle| / \sum_{hkl} \sum_j I_j(hkl)$, where N is the redundancy of the dataset.
[d]$CC_{1/2}$ is the correlation coefficient of the half datasets.

[e] $R_{work} = \sum_{hkl} \,||F_{obs}| - |F_{calc}|| \,/\, \sum_{hkl} |F_{obs}|$, where $F_{obs}$ and $F_{calc}$ is the observed and the calculated structure factor, respectively. $R_{free}$ is the cross-validation R factor for the test set of reflections (5% of the total) omitted in model refinement.

**Table S3:** Strains, plasmids, and primers used in this study.

| Strain, plasmid or primer | Characteristic(s) or sequence | Note |
|---|---|---|
| **Strains** | | |
| *L.lactis* (NZ3900) | MG1363 derivative | |
| **Plasmids** | | |
| pGIR210-LarAH31 | *Chl*$^r$. Production of LarA$_{lp}$ fused with a C-terminal Strep-tag | this study |
| pGIR210-Y294A | *Chl*$^r$. Production of the Y294A variant fused with a C-terminal Strep-tag | this study |
| **Primers (5'-3')** | | |
| LarAH31 Y294A-F | GATCTGACCTTCGCCCAAGCGGTGAAAG | mutagenesis |
| LarAH31 Y294A-R | CTTTCACCGCTTGGGCGAAGGTCAGATC | mutagenesis |
| LarAH31-SR | GTTGTAATATTTCTGCTGTGGTTGCC | sequencing |
| UP_PNZ8048' | ACAATGATTTCGTTCGAAGGAACTAC | sequencing |
